# ICTD: A semi-supervised cell type identification and deconvolution method for multi-omics data

**DOI:** 10.1101/426593

**Authors:** Wennan Chang, Changlin Wan, Xiaoyu Lu, Szu-wei Tu, Yifan Sun, Xinna Zhang, Yong Zang, Anru Zhang, Kun Huang, Yunlong Liu, Xiongbin Lu, Sha Cao, Chi Zhang

## Abstract

We developed a novel deconvolution method, namely **I**nference of **C**ell **T**ypes and **D**econvolution (ICTD) that addresses the fundamental issue of identifiability and robustness in current tissue data deconvolution problem. ICTD provides substantially new capabilities for omics data based characterization of a tissue microenvironment, including (1) maximizing the resolution in identifying resident cell and sub types that truly exists in a tissue, (2) identifying the most reliable marker genes for each cell type, which are tissue and data set specific, (3) handling the stability problem with co-linear cell types, (4) co-deconvoluting with available matched multi-omics data, and (5) inferring functional variations specific to one or several cell types. ICTD is empowered by (i) rigorously derived mathematical conditions of identifiable cell type and cell type specific functions in tissue transcriptomics data and (ii) a semi supervised approach to maximize the knowledge transfer of cell type and functional marker genes identified in single cell or bulk cell data in the analysis of tissue data, and (iii) a novel unsupervised approach to minimize the bias brought by training data. Application of ICTD on real and single cell simulated tissue data validated that the method has consistently good performance for tissue data coming from different species, tissue microenvironments, and experimental platforms. Other than the new capabilities, ICTD outperformed other state-of-the-art devolution methods on prediction accuracy, the resolution of identifiable cell, detection of unknown sub cell types, and assessment of cell type specific functions. The premise of ICTD also lies in characterizing cell-cell interactions and discovering cell types and prognostic markers that are predictive of clinical outcomes.

## Introduction

Tissue deconvolution aims to disentangle the cell composition in terms of their relative quantities, based on which, the cell type specific functions and their cross-talks in the tissue microenvironment could be studied ^1 2 3 4^. Existing deconvolution algorithms usually assume the observed expression matrix as a product of a cell type signature matrix S and proportion matrix P ^2 3 4^. Independent training data is usually needed to impose prior on S via certain information transfer ^2 5 6 7^. The recent emergence of single cell RNA-seq (scRNA-seq) allows researchers to uncover new biological traits in cell populations of bulk tissue^8^. Regardless, the knowledge transfer from training single/bulk cell data to target bulk tissue should be carefully handled, as the gene expression distribution of the two domains could be highly variable, which tend to be oversimplified in current deconvolution methods ^9^. Novel or rare cell subtypes are of great interest to researchers ^10^. However, current deconvolution algorithms usually assume a fixed pool of cell types, which clearly is incapable of identifying novel sub cell types ^2 3 4^. Moreover, certain cell types such as immune cells tend to co-infiltrate in a real tissue, suggesting that the proportions of these cell populations are highly co-linear ^11^. As a result, estimating proportions with plain linear regression model or non-negative factorization would suffer from multi-collinearity, leading to highly unstable predictions ^12 13^. Recent methods such as Cell Population Mapping (CPM) and CIBERSORTx have been developed to predict cell type specific functions ^14 9^. However, they rely on precisely predicted cell proportions, and matched scRNA-seq profiles of similar tissues, which limited their applications to a wider extent. It is also noteworthy that none of the existing deconvolution methods is designed to handle highly varied tissue microenvironments or multi-omics data. Here, we summarize the key challenges of deconvolution methods as (i) detect the resident (sub) cell types and their true marker genes dependent on the tissue ^15^ (ii) handle systematic expression variations from training to target data domain; (iii) deal with the prevalent co-linearity in the cell type specific expression signatures and cell proportions; (iv) define expression patterns that represent varied cell type specific functions; (v) enable application to a variety of tissue microenvironment and multi-omics data types. More detailed discussions and comparisons of the formulations of existing methods are provided in the **Supplementary Notes**.

Based on a preliminary evaluation of the variations of known cell type signature genes in a large set of single and bulk cell data, we first derived mathematical conditions for a cell type to be “identifiable” in a tissue omics data: (1) the cell type has uniquely expressed genes, the expression values of which over any subset of samples form a rank-1 matrix (a matrix with matrix rank equals to one), or (2) there are genes expressed by the cell type and other cell types satisfy (1), and the expression values contributed by the cell type over any subset of samples form a rank-1 matrix. And a cell type-specific function is “identifiable” if there are marker genes of the function forming a rank-1 submatrix in a subset of samples with significant presence of the cell type. These “identifiability” conditions grant the potential to detect novel cell subtypes or cell functions via the detection of low rank matrices. Detailed mathematical considerations and derivations were given in **Online Methods and Supplementary Notes**.

Based on the rigorously derived mathematical conditions, we developed a semi-supervised method, namely inference of cell types and deconvolution (ICTD), featured by: (1) a semi-supervised detection of “identifiable” cell types and marker genes specific to each omics dataset and tissue micro-environment; (2) a novel nonparametric detection and annotation of cell type signature genes, which is used as information basis to annotate the identified cell types; (3) a novel constrained non-negative matrix factorization (NMF) method to decrease the bias caused by knowledge transfer from training data, as well as to effectively handle the co-occurring cells; (4) a robust regression based approach to interactively deconvolute multi-omics data of matched samples, and (5) a local-low-rank screening approach to identify cell type specific functions, which altogether offers a systematic solution of the five key challenges.

## Results

Our core algorithm ICTD consists of six steps (**Fig 1**): <u>(1) Compute the relative specificity of all genes for all cell types in a given microenvironment.</u> A labeling matrix *L*_M×K_ of M genes and K selected cell types is first constructed based on training single or bulk cell transcriptomics data, where 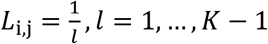, if gene *i* is significantly expressed in in cell type *j* and its expression is significantly lower than in *l* − 1 other cell types, and *L*_*i*,*j*_ = 0 otherwise (Supplementary Table S1). Without loss of generality, we assume that all the M genes are specific to one or a few cell types, namely, Σ_*j*_*L*_i,j_ > 0, ∀ *i* = 1, …, *M*. (2) <u>Detect all gene modules within which the gene expression vectors are linearly dependent and form rank-1 matrix, and the modules present evidence of “identifiable” cell types.</u> For each gene module detected on the target tissue expression matrix among the M genes, if its member genes are all highly expressed in one or several cell types according to the labeling matrix, the module will be considered as evidence of potential cell type(s) by ICTD. In this step, ICTD can exclude undesired cell types, such as cancer or other disease cells, from further analysis, by a non-negative projection of the input data to the complementary of the space spanned by the marker genes of undesired cell types. (3) <u>Infer the “identifiable” cell types and signature genes</u>. Non-negative linear dependency among the selected modules is evaluated and each module is annotated by the genes’ significant enrichment to a cell type based on the labeling matrix. Modules are merged with high inter-dependency, and further filtered such that modules enriching none of the cell types are removed. The total number of “identifiable” cell types is computed as the total rank of the expression matrix composed by all genes in the remaining rank-1 modules, the genes in each module will be considered as markers of the corresponding cell type. <u>(4) Predict cell proportions using constrained NMF.</u> With the “identifiable” cell types and their marker genes, a constraint matrix C_M×K_ can be constructed. Specifically, for cell type *k*, *k* = 1‥*K* with M_k_ marker genes, 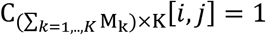, if gene *i* is marker of the cell type *j*, and 0 otherwise. The constraint matrix is then enforced upon the regular NMF formulation to guarantee similarity of the signature matrix with the constraint matrix, namely, we solve min 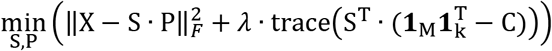, where **1**_d_ denotes an all-1 column vector of length *d*. (5) Co-deconvolution of matched multi-omics data. The semi-supervised property of ICTD enable its application to multi-omics data. A robust regression approach is further applied to identify the cell types and samples, in which the cell proportions inferred from different omics-data are highly consistent. <u>(6) Estimate cell type specific functions.</u> For each cell type detected, ICTD screens the rank of the expression matrix containing a group of samples which are stratified by their cell abundance levels, and pins down marker genes of a varied cell type specific function if they form at least one distinct dimension.

**Figure 1.**
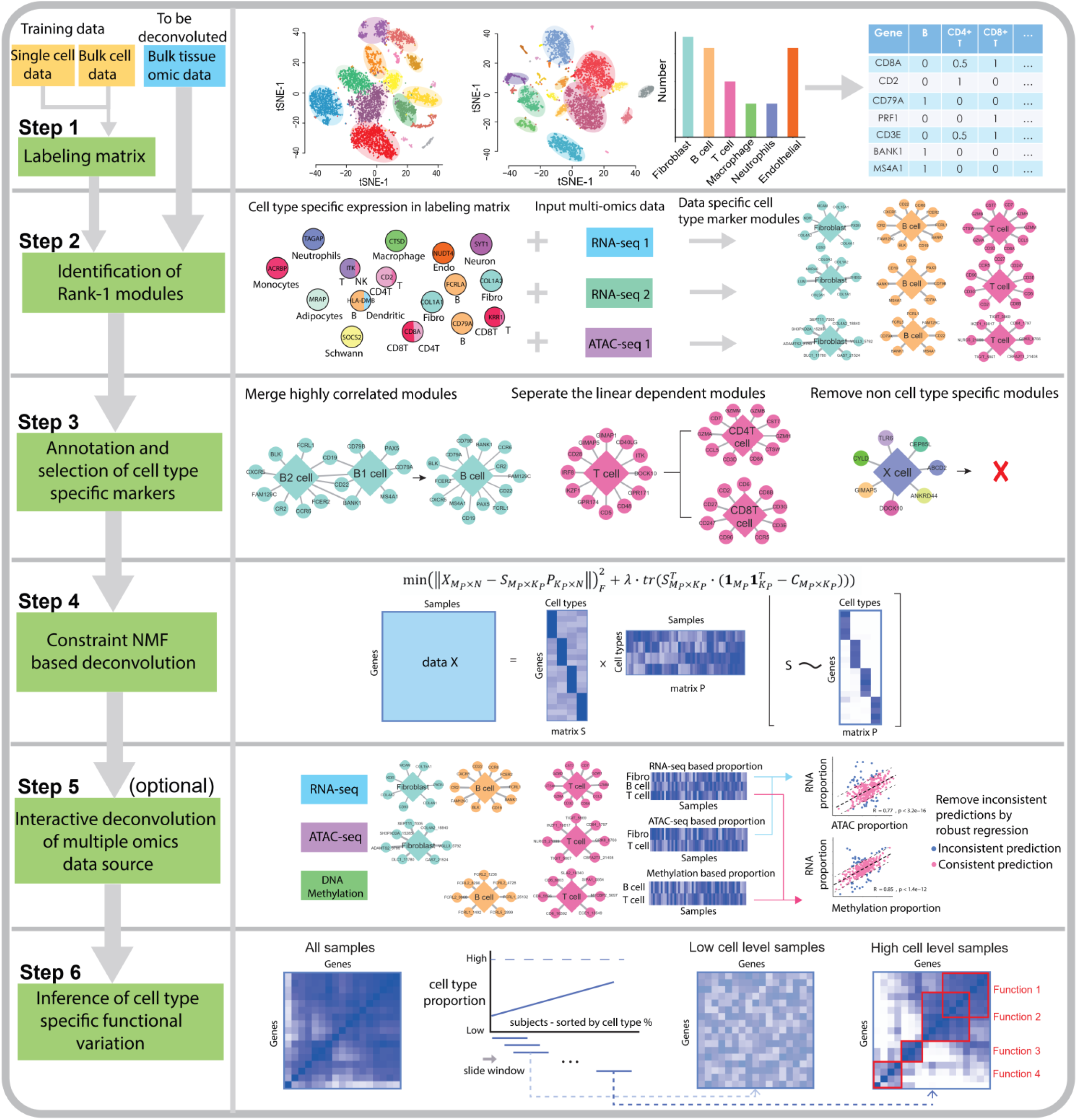
Analysis pipeline of ICTD. ICTD first constructs labeling matrix to store genes’ relative specificity to different cell types using bulk or single cell training data (Step 1). Rank-1 modules were detected among the cell type marker genes in each input omics dataset (Step 2). Similar modules were merged, modules that do not (non-negatively) depend on other modules are kept, and modules that do not overrepresent any cell type markers are removed. The number of cell types of the target deconvolution is determined as the total rank of the expression matrix of genes in the remaining modules (Step 3). A constrained NMF is conducted to regularize the signature matrix *S*, such that values in *S* are shrunken towards 0 if the corresponding entries in the constraint matrix is 0. (Step 4). If matched multi-omics data are available, robust regression among cell proportions inferred from different omics data set is performed to remove outlier samples (Step 5, optional). Marker genes of cell type specific functions are further identified by looking for local low rank submatrices in sample groups stratified by different level of the cell proportion (Step 6).

The core algorithms for each step are described in the **Online Methods**. Detailed algorithms, data used for method validation, and model comparisons with other methods, are provided in the **Supplementary Notes and Methods**. Below we present the application of ICTD on simulated bulk data using single cell RNA-seq data (**Fig. 2**) and real tissue data (**Fig. 3**). We demonstrated (1) the ability of ICTD to identify both known and novel (sub) cell types with high accuracy, (2) the overall competitive performance of ICTD in analyzing data of different tissue microenvironment and experimental platforms, (3) the robustness of ICTD in cases where cell types have highly co-linear proportions, (4) ICTD’s capability in interactive deconvolution of matched multi-omics data, (5) inference of cell type specific functions, and (6) explorative findings derived by correlating ICTD predicted cell and functional levels with other omics, imaging and clinical data.

**Figure 2.**
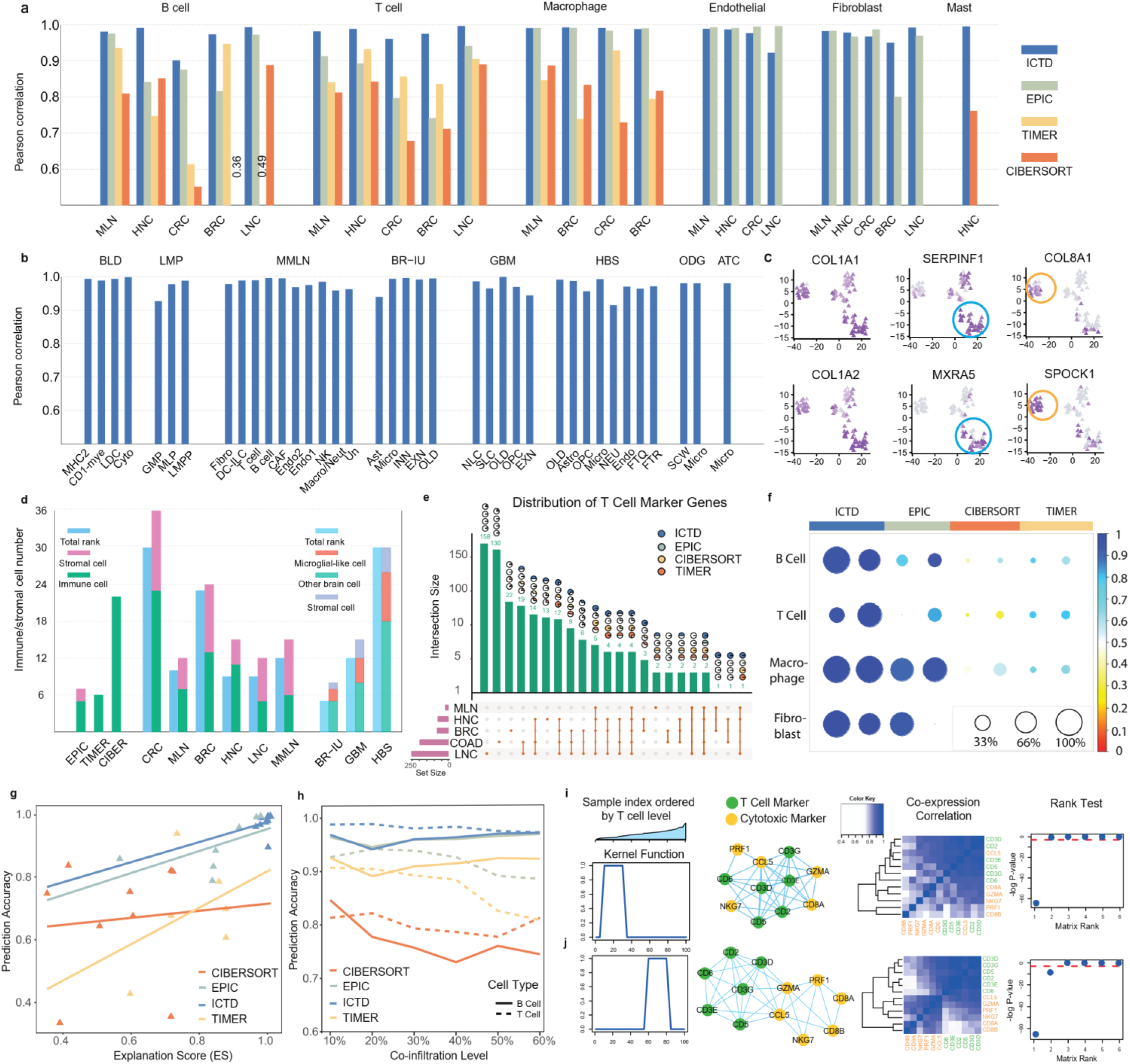
Validation of ICTD by using single cell simulated bulk tissue data. (a) PCC between true and predicted proportion of six cell types by ICTD, EPIC, TIMER and CIBERSORT, in the bulk tissue data simulated using scRNA-seq data collected from Melanoma (MLN), Head and Neck Cancer (HNC), Colorectal Cancer (CRC), Breast Cancer (BRC), and Lung Cancer (LNC). (b) PCC between true and predicted proportion of cell types and subtypes identified by ICTD in the bulk tissue data simulated by scRNA-seq data of myeloid and dendritic cell mixture (BLD), lymphoid and myeloid progenitor mixture (LMP), mouse melanoma (MMLN), normal brain cells nucleic sequencing generated in this study (BR-IU), glioblastoma (GBM), human normal brain (HBS), oligodendroglioma (ODG), and astrocytoma (ATC). Detailed cell type codes are given in **Supplementary Note**. (c) *t*-SNE plot of the marker genes of fibroblast subtypes in MLN scRNA-seq data, which were identified by ICTD from simulated human melanoma tissue data. In each panel, darker color denotes higher expression of the gene in a cell. (d) Consistency of the number of ICTD identified cell types and the matrix rank of the expression profile of the marker genes of identified cell types, i.e. the number of identifiable cell types, in each simulated tissue data. (e) Distribution of the true T cell marker genes identified in the five cancer data and their overlap with the actually used T cell signature genes in CIBERSORT, TIMER and EPIC. Each bar and number represent the number of genes specifically expressed by T cells in each of five cancer types, which is labeled in the dot plot on the bottom. The pie charts illustrate the proportion of the T cell marker genes used by ICTD (data adaptive) and CIBERSORT, TIMER and EPIC (held fixed). (f) Re-evaluation of robustness of cell type specific markers used by each method. The circle size represents the ratio of true marker genes among all genes used as marker genes for each cell type (row) for each method (column). The color represents the E-score level. The two columns of each method show the results of simulated MLN (left) and HNC (right) tissue data. The plots of the other three cancer types were shown in Supplementary Fig S6. (g) Dependency between explanation score (x-axis) and prediction accuracy (y-axis) of the cell type proportions given by the four methods. (h) PCC (y-axis) between true and predicted T and B cell proportions on simulated data with different level of T and B cell co-infiltration (x-axis). (i-j) prediction of varied T cell cytotoxicity level in simulated HNC data. From left to right, the four plots illustrate the kernel function used for local low rank screening, co-expression network of T cell and cytotoxic marker genes, heatmap of correlations between T cell and cytotoxic marker genes, and p values of the expression matrix rank of the T cell and cytotoxic marker genes, in the samples of low T cell infiltration (i) and high T cell infiltration level (j).

**Figure 3.**
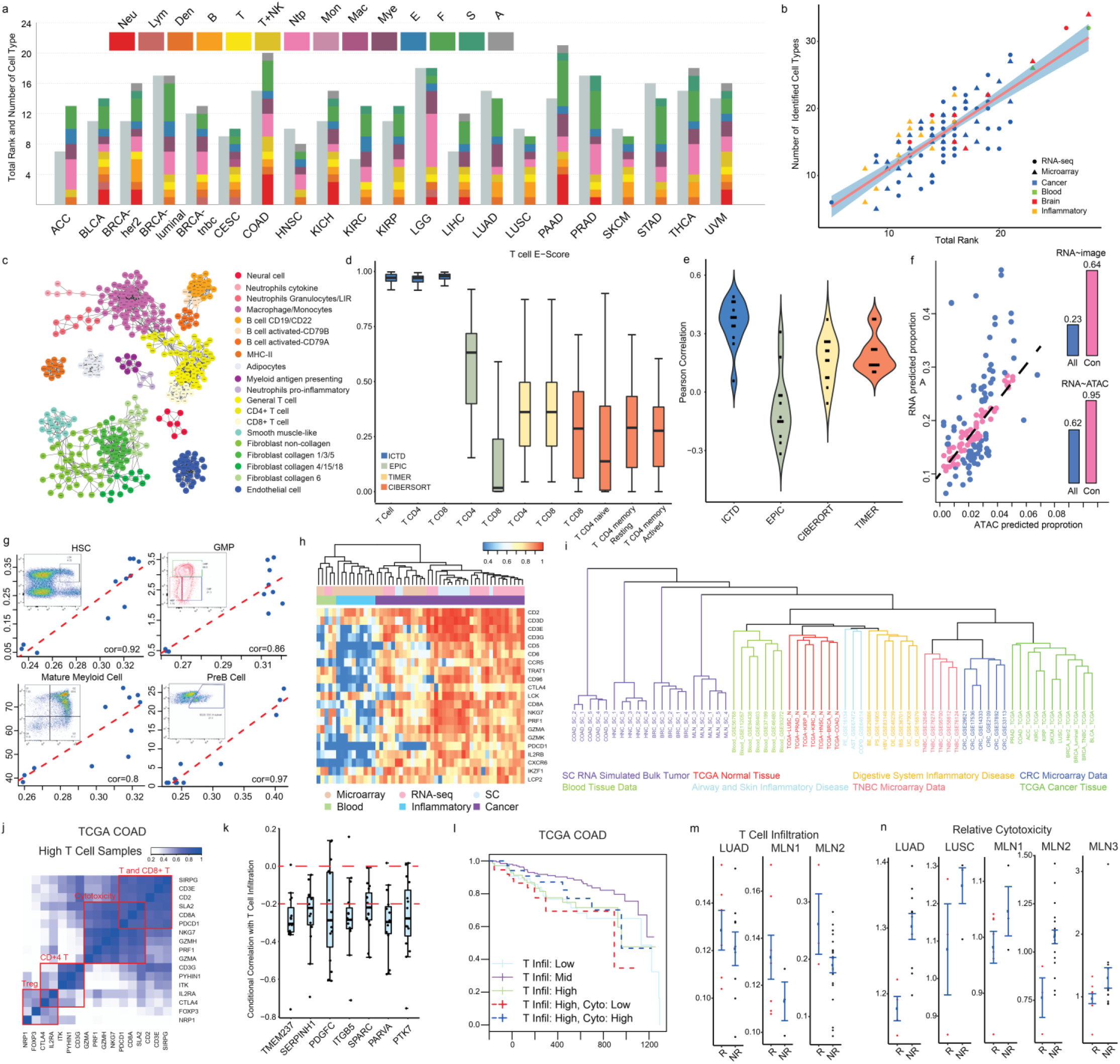
Application of ICTD on real bulk tissue transcriptomics data. (a) The number of identifiable cell types (colored bar) and the matrix rank (grey bar) of their marker genes identified by ICTD through all the TCGA cancer data; (b) Scatter plot of the number of identifiable cell types (y-axis) and the matrix rank (x-axis) of their marker genes identified by ICTD through all the analyzed data. (c) Network of the marker genes of the commonly identified cell (sub) types in the TCGA data. An edge between two genes means the two genes are both identified as markers of one cell type in more than 10 analyzed TCGA data. (d) E-score of the T cell marker genes identified by ICTD and those used by EPIC, TIMER and CIBERSORT in TCGA data. E-score of other cell types are given in Supplementary Fig S11. (e) Correlation (y-axis) between the imaging data derived tumor infiltrated lymphocyte level and T cell proportion predicted by the four methods (x-axis) in 11 TCGA cancer. (f) Scatter plot of the T cell proportions predicted by TCGA BRCA RNA-seq and ATAC-seq data. Samples with highest consistency identified by the robust regression were pink colored. The bar plots represent the correlations of the proportions inferred by the RNA-Seq vs ATAC-Seq (or RNA-Seq vs imaging) in all the samples (PCC=0.62, or 0.23) and the most consistent samples (PCC=0.96, or 0.64). (g) Consistency of ICTD predicted (x-axis) and FACS measured (y-axis) cell proportions of four hematopoietic cell types. (h) Evaluation of T cell markers identified in CNBI data. In the heatmap, each row is the commonly identified T cell markers and each column is one data set. Color in the heatmap represents the E-score of each gene in each data set. Statistics of other cell types are given in Supplementary Fig S12. (i) Clustering of datasets from different microenvironment from different platforms based on a distance measure of the marker gene expression profiles of identifiable cell types (see **Online Methods)**. This is to show the relative impact of technological platforms and tissue microenvironment on the variability of gene markers expressions. (j) Co-expression between T cell, CD8+ T cell, cytotoxic function, CD4+ T cell and T-reg marker genes in the samples with high T cell infiltration in TCGA COAD data. (k) Correlation between fibroblast cell expressing genes and T cell infiltration level conditional on the fibroblast cell level in 15 cancer types. (l) Survival curves of the TCGA COAD patients with low, medium and high T cell infiltration, and the high T cell infiltration patients with low and high cytotoxicity functions predicted by ICTD. (m) Variation of T cell infiltration level in response (R) and non-response (NR) patients in three independent checkpoint inhibitor treated clinical data. (n) Variation of T cell relative cytotoxic level in response (R) and non-response (NR) patients in five independent checkpoint inhibitor treated clinical data. LUAD, LUSC and MLN* represents different sets of lung adenocarcinoma, lung squamous cell carcinoma and melanoma.

### Validation on single cell simulated bulk tissue data

We benchmarked ICTD on predicting the types of resident cells and their relative proportions against three state-of-art deconvolution methods, namely CIBERSORT, TIMER, and EPIC (**Online Methods**), using single cell simulated bulk tissue datasets. The bulk tissue datasets were simulated by RNA-seq data of single cells or single nucleus from different tissue microenvironments, including five from human solid cancer (namely, breast, colon, head and neck, lung, and melanoma), five from human central nervous system (namely glioblastoma, oligodendroglioma, astrocytoma and two normal brain), three from human immune system (monocyte and dendritic cell, lymphoid, and myeloid progenitor cells), and one from mouse melanoma. On all five human solid cancer microenvironment, all mixing cell types were detected as “identifiable” by ICTD. In addition, ICTD achieved significantly higher accuracy in predicting total B-, T-, mast, fibroblast, endothelial cells and macrophage proportions comparing to other methods. On 23 out of the 25 cells type in the simulated bulk cancer datasets, ICTD predicted relative proportions achieved higher than 0.95 Pearson correlation coefficient (PCC) with true proportions, while the average PCC is 0.86, 0.63 and 0.52 for EPIC, TIMER and CIBERSORT, respectively (**Fig 2a**). On the five human brain microenvironments, ICTD successfully detected astrocyte, oligodendrocyte and progenitors, exhibitory and inhibitory neuron, microglial and Schwann cells as identifiable cell types, all with at least 0.9 PCC with true proportions (**Fig 2b**). Similarly, ICTD also accurately identified sub ell types from the mixture of multiple classes of monocyte and dendritic cells, human lymphoid and myeloid progenitors, and the immune and stromal cells in mouse melanoma microenvironment, with reliable prediction of proportions (**Fig 2b**).

#### Novel cell types

A unique feature of ICTD is its capability to automatically detect cell and sub cell types along with cell marker genes for effective cell (sub)type annotation. Our analysis on simulated cancer tissue data suggested that each of the rank-1 module corresponds to one cell or sub-cell type (Supplementary Fig S1). On the simulated human solid cancer datasets, ICTD was able to identify subtypes of immune/stromal cells, including CD4+ and CD8+ T cells, novel subtypes of fibroblast and myeloid cells. The sub cell type markers were further validated by the tSNE visualization, where the expression level of each marker set turns out to be specific to the associated cell types or subtypes (Supplementary Fig S2). As illustrated in **Fig 2c**, among the three fibroblast rank-1 modules identified by ICTD, one clearly corresponds to the general fibroblast (with COL1A1 expression) type, and the other two correspond to two fibroblast subtypes (with COL8A1 or SERPINF1 expression) in the simulated human melanoma data. We confirmed all the rank-1 modules identified by ICTD from all the single cell simulated tissue data are specifically expressed in only one cell type, suggesting the high specificity of ICTD in identifying true cell types.

#### Variability of cell types and their marker genes

It is noteworthy that the number of identifiable cell types could vary through disease contexts and data sets. Comparing to the fixed cell types assumed in most of the deconvolution methods, the number of cell types identified by ICTD highly matches the number of mixing cell types in each single cell simulated tissue data set (**Fig 2d**). We further investigated the level of variation for cell type markers through different disease contexts and data set. As shown in **Fig 2e**, there is a strong disease context specificity of T cell markers: only four T cell markers were shared by all the five cancer data sets, and 19 T cell markers were shared in four out of the five data sets. We observed on average 93.75%, 90.36% and 83.33% of the T cell markers utilized in CIBERSORT, TIMER and EPIC are specific to only three or less cancer types and only 65.21%, 69.57% and 13.04% of the common cell type marker genes were included in their signature matrix. Similar patterns are also seen for B and fibroblast cells (Supplementary Fig S3). In contrast, ICTD considers the variations in both the identifiable cell types and cell type markers in different tissues and datasets, resulting in a better prediction accuracy throughout different scenarios (**Fig 2e-f**). An explanation score (ES) is defined for each marker gene to evaluate the goodness of fitting of the gene’s expression by the predicted proportions of the cell types expressing the gene (**Online Methods**). High ES scores of the marker genes for one cell type is a necessary condition for the high prediction accuracy and specificity of the marker genes. We observed strong positive correlations between the ES scores and prediction accuracy using ICTD and EPIC, as these two methods rely on cell type uniquely expressed genes. Similarly, for CIBERSORT and TIMER, positive associations were also observed (**Fig 2g**). Analysis of six major immune and stromal cell types in five simulated bulk cancer data sets suggested that in general, when ES is below 0.8, the prediction accuracy is lower than 0.8; on the other hand, when ES is above 0.9, the prediction accuracy tends to higher than 0.9 (**Fig 2g**). We observed the ES of all the cell type specific markers identified by ICTD on the simulated cancer tissue data are all above 0.95. It is noteworthy ES can partially evaluate the performance of a deconvolution method without knowing true cell proportions.

#### Cell type co-linearity

ICTD also demonstrated its superiority in handling co-linearity of cell proportions, caused by cells’ functional dependencies. Our preliminary analysis on TCGA data suggested correlation among the immune and stromal cells to be as high as 0.94 (**Supplementary Notes**). We simulated batches of bulk tissue samples in each of which the cell proportions are intentionally set to have different levels of correlations to mimic the dependencies of different cell types in cancer microenvironment (**Online Methods**). Not surprisingly, while performance of regression based methods dropped significantly when co-linearity level was high, ICTD achieved high robustness and prediction accuracy at different levels of co-linearity. This owes to the data-adaptive selection of cell type specific markers and constrained NMF formulation adopted by ICTD. The four methods’ prediction accuracy of B and T cells across different co-linearity levels in simulated human melanoma tissue data is shown in **Fig 2h**. In addition, significant correlations among ES, prediction accuracy, and co-linearity of cell proportions were identified (Supplementary Fig S4).

#### Cell type specific functions

ICTD can identify varied function of a certain cell type using a local low rank identification approach ^16^. In the human head and neck cancer data, we identified the expression level of cytotoxic gene in the CD8+ T cells vary considerably in patient stratifications of different T cell abundances, suggesting mixed T cell exhaustion levels (**Supplementary Notes**). To evaluate the capability of ICTD in identifying varied T cell cytotoxicity level, we simulated bulk tissue data with different proportion and cytotoxicity level of T cells (**Online Methods**). ICTD conducted a local low rank screening with a kernel function along samples ordered by predicted T cell proportions. Our analysis clearly identified the linear space spanned by the T cell and cytotoxicity marker genes switches from rank-1 to rank-2 throughout the samples with low to high T cell levels, suggesting the identifiability of the varied cytotoxic level in the samples of high T cell infiltrations (**Fig 2i-j**). On average, correlation level of 0.86 between the true cytotoxicity level per unit T cell and the prediction made by ICTD was observed (Supplementary Fig S5).

### Implications from real tissue data

We then applied ICTD on a collection of human cancer, normal, blood and inflammatory tissue (CNBI) data, including 28 cancer and 11 normal tissue types from TCGA, 17 colorectal cancer, 7 triple negative breast cancer, 7 blood tissue, and 11 human inflammatory disease data sets from GEO (Supplementary Table S3). We identified rank-1 markers of B, T, dendritic, general myeloid, macrophage, monocytes, neutrophil, fibroblast, endothelial and adipocyte cell and their sub cell types in each dataset (**Fig 3a** and Supplementary Fig S7). A strong association between the number of identified cell types and the total rank of the matrix of marker genes was observed (**Fig 3b**). It is noteworthy that the types of resident cells most variable across different cancer types are the subtypes of adipocytes, fibroblast, and myeloid cells, which seem to be most prevalent in breast, colorectal, lung, pancreatic and stomach cancers, commonly known to have considerable stromal components. The complete set of cell types and their marker genes identified in each data set were summarized in Supplementary Table S4. In the TCGA datasets, 21 commonly “identifiable” cell and subtype types have been observed in more than 10 cancer types, including CD19/CD22 expressing regulatory-like B cell and CD79A/CD79B expressing activated B cell; total, CD8+, and CD4+ T cell; Neurexin and Caytaxin expressing Neuron cell; myofibroblast-like cell; Collagen 1/3/5, Collagen 4/15/18, Collagen 6, and Non-collagen expressing Fibroblast; Endothelial cell; MHC class II antigen presenting cell; MHC class I, pro-inflammatory cytokine releasing, chemokine and cytokine releasing Myeloid cells; complement pathway activated Macrophage and Monocytes; granulocytes; and adipocytes (**Fig 3c**).

We confirmed markers of each commonly identifiable cell types in cancer microenvironment do have significant overlaps with the immune and stromal cell markers identified in normal microenvironment, suggesting these marker genes truly belong to immune and stromal cells rather than cancer cells (Supplementary Table S5). On average, the ICTD marker genes of each cell type have ES higher than 0.9, while the ES scores of the signature genes used by CIBERSORT, TIMER and EPIC are 0.22, 0.39 and 0.26, respectively. **Fig 3d** illustrate the ES of T cell (sub)type markers of the four methods. The level of tumor infiltrated lymphocytes (TIL) in 12 TCGA cancer types have been previously assessed by imaging data ^17^. On average, the correlation between imaging predicted TIL and ICTD predicted T cell level is 0.4, comparing to 0.14, 0.2, and −0.11 with CIBERORT, TIMER, and EPIC predicted T cell level (**Fig 3e**). For other cell types, with a lack of ground truth, we rely on evaluating the ES scores of 3,552 known immune and stromal cells marker genes. It turns out that ICTD-predicted cell proportions achieved on average 0.56 R^2^ value in explaining the 3,552 known immune and stromal cells marker genes, while the R^2^ is 0.2, 0.24, and 0.18 for CIBERSORT, TIMER, and EPIC (Supplementary Table S6).

ICTD enables interactive deconvolution of matched multi-omics data. We co-deconvoluted the RNA-seq, ATAC-seq and DNA methylation data of five TCGA cancer types with available data (Online Methods). On average, more than 70% of the cell types identified from RNA-seq data were also identified in ATAC-seq or methylation data, including adipocytes, B cell, CD4+ and CD8+ T cell, macrophage, fibroblast, endothelial and dendritic cells (Supplementary Table S7). The correlations between cell proportions inferred from different data types are higher than 0.6. Fig 3f illustrated the strong consistency between the T cell proportion inferred from TCGA BRCA RNA-seq and ATAC-seq data. It is noteworthy the samples used in multi-omics experiments were from different parts of a tumor tissue, and some are less representative of the whole tumor tissue. ICTD utilizes a robust regression approach to remove such samples with inconsistent cell proportions inferred from the multiple data sources. As a result, the correlation between RNA-seq and imaging inferred T cell proportion was increased from 0.23 to 0.64, wherein the imaging based proportion is deemed as a reliable reference here. This suggests the interactive co-deconvolution of multi-omics data has the potential to increase the robustness of the prediction.

Application of ICTD on 7 human normal brain, 5 neuro-degenerative disease and 4 brain cancer data sets identified 23 common cell types in central nervous system, including two astrocyte, three general glial, two oligodendrocyte, oligodendrocyte progenitor, exhibitory and inhibitory neuron, MHC class I and II antigen presenting cells, general myeloid, macrophage, neutrophil and stromal like microglial cells, one endothelial, one epithelial, three ependymal, and one collagen expressing stromal like cell types (Supplementary Fig S8).

To experimentally validate ICTD in identifying rare sub cell types and predicting cell proportions in complex tissue system, we generated an RNA-seq data set of 12 mouse bone marrow tissue samples each with flow cytometry (FACS) measured cell numbers (see details in **Supplementary Notes**). ICTD successfully identified all the four hematopoietic cell types measured by FACS, namely hematopoietic stem cell, general myeloid progenitor, mature myeloid cell and pre-B cell, and achieved correlations of 0.92, 0.86, 0.8 and 0.97 between predicted and FACS measured cell proportions. Complete statistics including labeling matrix of mouse hematopoietic cell types, cell type specific markers identified by ICTD, cell proportions predicted by ICTD and measured by FACS were given in **Fig 3g**, Supplementary Table S8 and Supplementary Fig S9.

ICTD considers the variability of resident cell types and their marker genes across tissue microenvironments and technology platforms. **Fig 3h** illustrate the ES of T cell expressing genes in different CNBI data sets, suggesting a significant variation of the T cell markers in the microenvironment of different cancer, inflammatory disease and blood tissue, as well as under different experimental platforms ^18 19^. To further investigate how the data set specific makers vary by disease/tissue micro-environments or experimental platforms, we further computed the averaged Jaccard distance between the marker genes of same cell types identified in any two CNBI or single cell simulated bulk datasets (**Supplementary Methods**). As illustrated in **Fig 3i**, the cell type marker genes vary drastically between cancer, normal inflammatory and blood tissues. Three distinct clusters were observed (1) TCGA cancer and other cancer, (2) single cell simulated cancer, and (3) TCGA normal and other inflammatory disease, and blood tissue. Among the cancer data, TCGA and other RNA-seq based cancer data sets is well separated from scRNA-seq simulated cancer data and the Microarray cancer data sets, and the later one is further divided into two sub-clusters containing independent CRC and TNBC data sets. Similarly, the TCGA RNA-seq and microarray data of normal, inflammatory conditions, and blood tissue form three distinct sub-clusters. Among the microarray data of chronic inflammatory conditions, the disease of digestive system and airway and skin tissues from two sub-clusters.

ICTD detected general T cell, fibroblast, and myeloid cells in all 28 analyzed TCGA cancer types, while the CD8+ T, non-collagen extracellular component expressing fibroblast, and oxidative stress producing myeloid cells were identified as distinct cell types in only 10, 12, and 15 cancer types, respectively. We found that the markers of these functional sub cell types are detected as cell type specific functions instead of a cell type in some cancer types by the local low rank screening function. For the 19 cancer types where CD8+ T cell is not identified as a cell type, CD8+ T cell markers were treated as one T cell specific function in 15 cancer types, while in 4 cancer types, high concordance is observed between total T cell and CD8+T cell markers in all the samples, making the CD8+ T subtype not differentiable from the general T cell. **Fig 3j** illustrated the marker genes of general T, CD8+ T, CD4+ T and T-reg cells form a distinct rank-4 submatrix in samples with high T cell infiltration, while the genes were less distinguishable in the complete TCGA COAD data (Supplementary Fig 10). This suggests the “locality” of finding identifiable cell types and functions, and hence it is necessary to implement a local low rank module detection approach. Similar locality was also observed for the marker genes of non-collagen expressing fibroblast and NADPH oxidase expressing myeloid cells in certain TCGA cancer types and other analyzed CRC and TNBC data sets (Supplementary Fig S10). We also conducted comprehensive screening to identify unknown immune/stromal cell type specific functional genes (**Online Methods**). 84 major functional modules were identified as common cell type specific functions in TCGA data (**Supplementary Notes**).

#### Cell-cell interaction

The prediction of cell proportions and functions by ICTD makes it possible to computationally characterize cell-cell interactions. We observed co-infiltrations among immune and stromal cell types with PCC in the range of −0.2-0.94 in all the analyzed TCGA cancer data (Supplementary Table S9). More importantly, the functional promotion or inhibition of cell type A to cell type B could now be examined by the correlations between the abundance level of A and the activity level of the function in B, conditional on the predicted proportion of B. We found seven genes expressed by fibroblast cells with significant negative conditional correlation with T cell infiltration in at least 10 out of 15 cancer types with high level of stromal cells (p<0.01) (**Fig 3k**). The seven genes execute functions related to the modification and synthesis of collagen and extracellular polysaccharide, suggesting a possible role of the dysregulated extracellular matrix composition in directing T cell infiltration. Similarly, the interactions of functions in two cell types can be computed by the correlation of the activity levels of the two functions conditional to their proportions. We identified a low conditional correlation among CD8 T cell markers such as CD8A/CD8B and cytotoxic genes, and a high conditional correlation among general T, CD8+ T, and cytotoxic genes in 4 cancer types, suggesting possibly perturbed cytotoxicity of T cells in the first 19 cancer types, namely T cell exhaustion. We also observed a significant negative correlation (p <0.01) between the NADPH oxidase and T cell cytotoxicity levels conditional to the total myeloid and T cell in 11 out of the 25 TCGA cancer types (Supplementary Table S10). This is consistent with previous observation that NADPH oxidases produce reactive oxygen species (ROS) on the surface of myeloid-derived suppressor cells that suppress the cytotoxic function of T cells ^20^.

#### Clinical implications

ICTD enables investigation of the impact on clinical prognosis by microenvironment. We conducted association analysis between the predicted cell proportions and varied functions with patient’s overall survival in TCGA data, as well as patients’ response in five clinical trial data with immune checkpoint inhibitor treatment (**Supplementary Methods**). We identified significant associations of patients’ overall survival with T cell infiltration and relative cytotoxicity levels in 12 and 7 TCGA cancer types, respectively. More interestingly, in colorectal and ovarian cancer, we observed that patients with moderate level of T cell infiltration have the best overall survival comparing to the patients with high and low T cell levels (**Fig 3l**). We define the T cell’s relative cytotoxicity (RC) level as the predicted cytotoxic function. level divided by the predicted total T cell level in each sample and observed patients with higher RC have significantly better overall survival. This clearly suggests the existence of T cell exhaustion and its association with poor prognosis. On the five clinical trial data, we noticed that patients with high T cell infiltration have better response to the treatment (**Fig 3m**), which is consistent with previously reported ^21^. Moreover, the level of T cell cytotoxicity was observed to vary significantly in four datasets of melanoma, lung adenocarcinoma and lung squamous carcinoma. We observed the patients with lower RC tend to have better clinical response (**Fig 3n**), possibly due to more PD-1/PD-L1-mediated immuno-suppression in these tumors. It is noteworthy that association between T cell infiltration and patients’ clinical outcome, and the identifiability of varied cytotoxic function show a high consistency between TCGA and the clinical trial data (Supplementary Table S10).

## Discussion

Our semi-supervised deconvolution method ICTD brought up a novel notion called “identifiability” of a cell type and cell type specific function, which was mathematically rigorously defined. By adaptively defining detectable cell types and selecting cell type markers based on the input data resolution, ICTD highly reduces the estimation bias, and also enables detection of novel cell (sub) types, and cell type functional activities. These features are particularly favorable when the goal is to computationally characterize the cell-cell interactions in large-scale tissue transcriptomic profiles. It is noteworthy that the “transcriptionally identifiable” cell types differ from those defined by cell differentiation lineage: some cell types on the lineage map may not be identifiable, while an “identifiable” cell type can be a certain cell or cell subtype, or the total of several cell types on the lineage map that express same gene markers. We believe the liberty of ICTD in its deconvoluted cell types makes it entirely data-driven, less biased to the training data, and it thus grants more sensible findings for downstream correlation analysis with other clinical and biological features.

ICTD is flexible in utilizing different types of training data to construct the labeling matrix, and we noticed using scRNA-seq profiles of cells from the real microenvironment of a certain cancer type, we are able to derive more tissue specific cell type markers than using microarray expression profiles of primary cells collected from healthy donors (**Supplementary Notes**). It is also worthy of mention that since ICTD is not fully supervised, we suggest at least 10 samples is needed for the method to work. While the method has increased type II error when the sample size is small, the identified rank-1 gene modules can be informative in guiding the flexible selection of cell type signature genes. Based on this, our ICTD R package was integrated with a regression based approach specifically for small sized samples with data-guided gene markers. When multi-omics data is available, we showed that co-deconvolution of matched multi-omics data could improve the prediction robustness, by excluding certain “outlier” samples with unstably predicted proportions using robust regression, and this function is available in the ICTD R package. The R package and web server version of ICTD are available at https://github.com/changwn/ICTD and https://shiny.ph.iu.edu/ICTD/.

Application of ICTD on TCGA pan-cancer data identified variations of T cell marker, cytotoxic marker and T cell exhaustion level, association between fibroblast expressing genes and T cell infiltration level, and association between ROS produced by myeloid cell and T cell cytotoxic level in different cancer types, suggesting the capacity of ICTD in providing a comprehensive evaluation of tissue specific cell types, cell type specific function, and cell-cell interactions. Nevertheless, the sensitivity of detecting cell type varied function can be largely improved if more prior knowledge of functional marker genes is available. And additionally, more novel cell type functions can be predicted if the rank-1 module detection approach could be optimized such that certain modules may exist with respect to only a subset of samples, considering the prevalence of disease heterogeneity and subtype specificity. In other words, co-expression modules local to subset of samples may be desirable in revealing more cell type functions.

## Online Methods

### Single cell, bulk cell and tissue transcriptomics data sets used in this study

We collected bulk cell data of 11 types in human blood, inflammatory and cancer tissue microenvironment, 8 types in human central nervous system, all generated by Affymetrix UA133 plus 2.0 Array; and 13 types in mouse inflammatory and tissue microenvironment, generated by Affymetrix Mouse Genome 430 2.0 Array. Detailed cell types include: *human stromal and immune cells:* fibroblast (34, 387), adipocytes (3, 26), endothelial cell (29, 606), B cell (20, 404), CD4+ T cell (23, 443), CD8+ T cell (9, 130), natural killer cell (9, 141), dendritic cell (32, 410), monocytes (22, 477), macrophages (21, 277), and neutrophil (10, 257); *human central nervous system:* neuron (16, 243), Schwann cell (2, 14), astrocyte (10, 57), ependymal cell (1, 39), oligodendrocyte (4, 30), and microglial cells (43,754), endothelial (29, 606), and stromal-like cell (34, 387); *mouse stromal and immune cells:* fibroblast (28, 277), adipocytes (3, 63), myocytes (myocyte), endothelial cell (10, 56), B cell (6, 31), CD4+ T cell (6, 80), CD8+ T cell (3, 34), natural killer cell (7, 35), dendritic cell (12, 84), monocytes (10, 46), macrophages (8, 102), neutrophils (11, 36), and mast cell (3, 31). The two numbers in the parenthesis indicate the number of datasets and samples of each cell type. We believe these cell types, together with tissue primary cells can cover major cell populations in the microenvironment of solid cancer, inflammatory disease, central nervous and hematopoietic system. 2854 samples of cancer cell line, human and mouse tissue index, and other cancer and normal tissue data were utilized as background to exclude the genes expressed by cancer or tissue primary cells.

The method was validated on single cell simulated bulk data. 13 single cell RNA-seq data sets generated by either C1/SMART-seq2 or 10x Genomics pipelines are used, and the cells are collected from (1) the TME of human solid cancer melanoma (8, 4486), breast (7, 535), colorectal (8, 375), head and neck (9, 5902), and lung cancer (8, 6630), (2) human glioma (5, 751), oligodendroglioma (7, 2728), and astrocytoma (7, 5171), (3) one public (8, 420) and one in-house (5, 1239) human normal brain sets, (4) human myeloid cell lineage and lymphoid cell lineage (3, 318) and monocyte/dendritic cell populations (4, 700), and (5) the TME of mouse melanoma (9, 2903). The two numbers indicate the number of cell types and cells of each data set.

We applied ICTD on real bulk tissue transcriptomic data of (1) 28 TCGA cancer types, (2) 11 TCGA normal tissue data, (3) 17 independent microarray data sets of colorectal cancer measured by different platforms; (4) metabric and 6 other triple negative breast cancer data sets; (5) 7 blood tissue RNA-seq and microarray data; 11 human inflammatory disease data sets generated by Affymetrix UA133 plus 2.0 Array, and (7) 7 human normal brain, 5 neuro-degenerative disease and 4 brain cancer types. Detailed information of the bulk cell, scRNA-seq and bulk tissue data were provided in Supplementary Table S3. The sample information and selection, downloading and processing procedures of the public data, and sample and sequencing information of the inhouse generated data were given in **Supplementary Notes**.

### Preliminary derivation of the mathematical conditions of “Identifiable” cell types and cell type specific functions

As detailed in **Supplementary Notes**, we analyzed the following characteristics of the cell type signature genes in the scRNA-seq and bulk tissue data of different disease context, experimental platforms and batches: (1) the consistency of cell type uniquely expressed genes were evaluated by their averaged expression level in different cell types of different scRNA-seq data sets; (2) inter- and intra-sample variations of cell type signature genes were characterized by the “drop-out” rates and multimodality of each gene’s expression profile in the scRNA-seq data of different samples; (3) matrix rank and expression scale of cell type uniquely expressed genes in bulk tissue data were evaluated by using BCV based rank test and Kolmogorov Smirnov (KS) test, and (4) immune and stromal cell co-infiltrations in cancer and inflammatory tissues were further assessed by using the averaged co-expression correlations among a small number of known cell type uniquely expressed genes.

Our evaluation suggested that NMF solution may not be unique if the used marker gene set are expressed by more than one cell type due to the prevalent co-linearity of cell proportions (**Supplementary Notes**). Hence only the cell type with uniquely expressed genes are transcriptomically “identifiable”, and the markers genes should also be stably expressed through cells of the same type so that its tissue level expression can reflect the cell’s population in the tissue. Specifically, if gene *i* is uniquely and stably expressed in cell type *k*, its gene expression can be expressed as 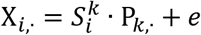, where 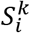 is the unit expression of *i* in *k*, and 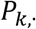 is the relative proportion of cell type *k* across all the samples. This shows that genes uniquely expressed by a cell type forms a (matrix) rank-1 submatrix, which form a necessary condition of “transcriptomically identifiable” cell type. On the other hand, a significant rank-1 structure of the expression profile of multiple genes 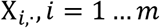 suggests that these genes are highly possibly expressed by a dominating cell type in the current tissue microenvironment or the genes are with similar expression pattern in several cell types.

Noting cell type specific functional activities, such as the T cell cytotoxicity, are highly varied through different patients, it is not feasible to use constant gene expressions level to characterize their activities. Denote the averaged level of a functional gene *i* in cell type k in the sample j as 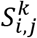, our evaluation suggested that the function is identifiable only if there exists a group of marker genes *i* = 1 … *K* satisfy 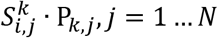 form a rank-1 matrix. Specifically, the cell type specific functional genes should share the same rank-1 space with the cell type markers if there is no variation while the functional genes can be identified as the markers of a cell type if 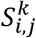 varied in all samples. If only a subset of samples has the functional variation, the low rank structure of the functional genes will be absorbed by the cell type markers and diminish on the co-expression network of all the samples. For such a case, the linear base of the varied function can be distinguished when the computation was limited to the samples with the functional variation, i.e. a local low rank identification method is needed (See more discussions in **Supplementary Notes**).

### A modified Bi-cross validation (BCV) based test of matrix rank

Bi-cross validation (BCV) has been developed to estimate the matrix rank for singular value decomposition (SVD) and Non-negative Matrix Factorization (NMF), which requires a prefixed low dimension *K* and two low rank matrices for the approximation *X*_*M*×*N*_ = *W*_*M*×*K*_ · *H*_*K*×*N*_.. The error distribution of gene expression data is usually non-identical/independent, majorly because a gene’s expression can be affected by its major transcriptional regulators, other biological pathways and experimental bias. Hence undesired biological characteristics and experimental bias may form significant dimensions in a gene expression data ^22^. In sight of this, we developed a modified BCV rank test (**Algorithm 1**) to minimize the effect of the non-i.i.d errors in assessing the matrix rank of a gene expression data.

**Algorithm 1:**
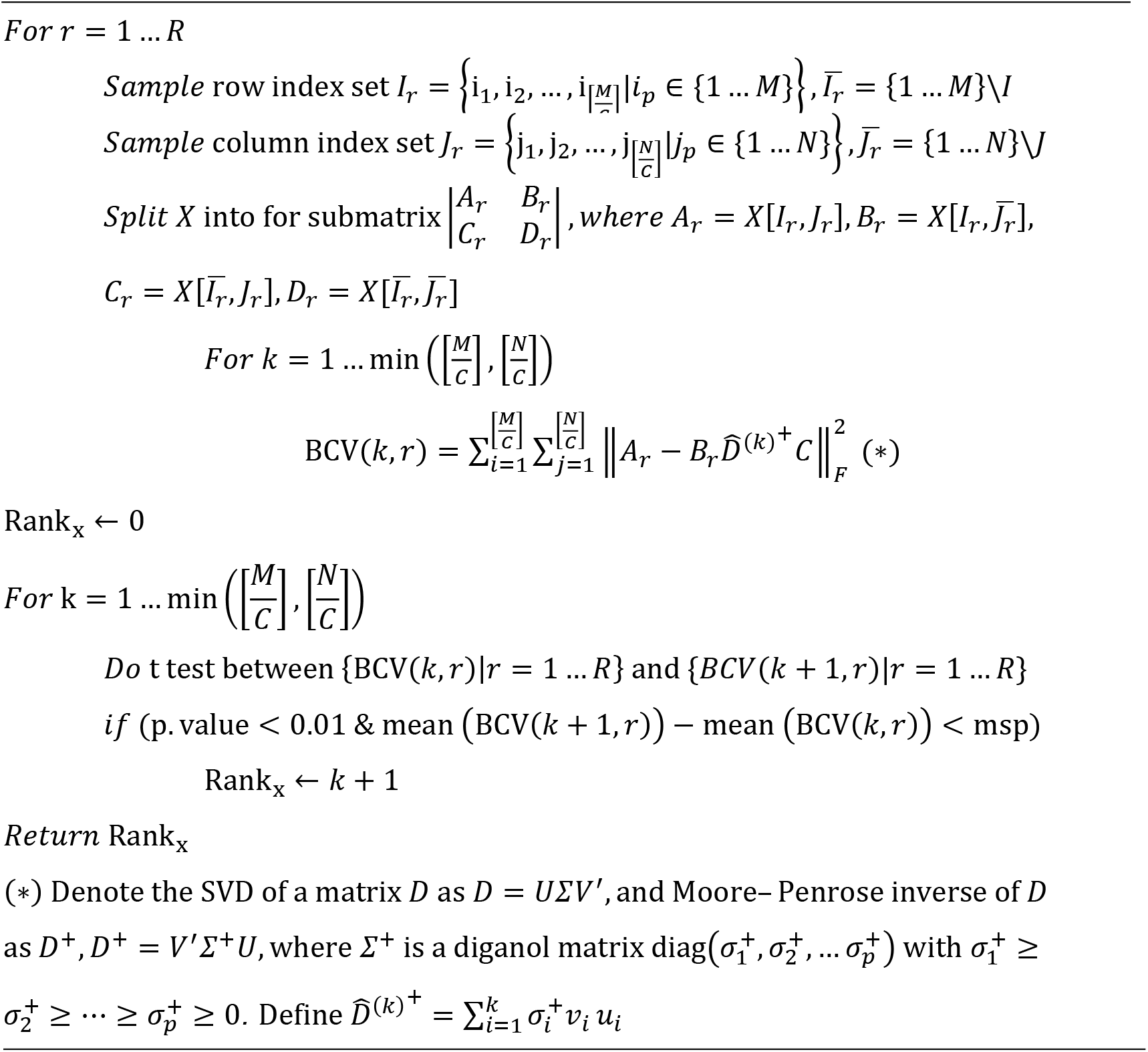
Modified Bi-cross validation matrix rank test

### ICTD Step 1: Construction of labeling matrix to represent TME specific cell type marker genes

A labeling matrix *L*_*M*×*K*_ was first constructed to represent the genes that are overly expressed in a certain cell type, where *M* is the number of genes and *K* is the number of cell types, 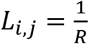 stands for the gene 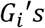 expression in cell type *C*_*j*_ is the *R*th highest among its expression in all the cells, and *L*_*i*,*j*_ = 0 stands for *G*_*i*_ is not a significant signature of cell type *C*_*j*_. Two different approaches were developed to construct the labeling matrix by using scRNA-seq or bulk cell data:

1. scRNA-seq data: For a scRNA-seq data set with annotated cell labels of *K* cell types and a given gene *g*, denote the expression profile of *g* in cell type k as 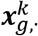, its mean as 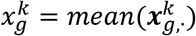, and the Z score of 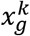 as 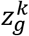. The cell type order vector ***o*** was further computed, where ***o***_*j*_ = *k*, if the *j*th largest value of 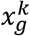 happens to be of cell type *k*. Then for cell type ***o***_1_ to ***o***_***K***_, the labeling matrix was built by

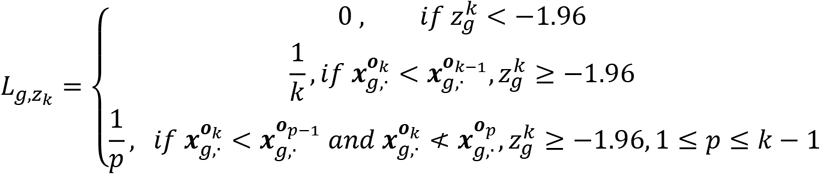

where 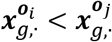 denotes *g* is significant over expressed in cell type ***o***_*j*_ compare to cell type ***o***_*i*_, which is tested by using MAST ^23^.
2. bulk cell data: We applied a non-parametric random walk based approach to identify if a gene has higher expression in certain cell types comparing to others, i.e. a signature gene of the cell types, by using the training data set composed by a large independent data sets of the cell types. ICTD enables the user to select the cell types specific to a tissue microenvironment. For examples, bulk cell data of normal breast cell, breast cancer cell lines and breast cancer tissue samples were selected as background to train the marker genes of immune and stromal cells for analyzing breast cancer tissue data. The labeling matrix used in this paper were computed by using human CCLE cell line, human body index and more than 20 human cancer tissue data as the background data. Batch effect of the training data of each cell type were first removed by using COMBAT ^24^ and the expression profile of each sample was further normalized by its mean. Denote the combined expression matrix containing *M* genes for *N* samples of *K* cell types, and for each cell type, we first calculated the expected frequency of the cell type, i.e. dividing the total number of samples for the cell type (*N*_*k*_, *k* = 1, …, *K*) by the total number of samples *N*, denoted by *E*_*i*_ = *N*_*k*_/*N*, *i* = 1, …, *K*. For a given gene *g*, denote 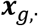 **and** 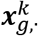 as its expression profile of all cell types and cell type *k*. We order the corresponding cell type labels of these samples based on the expression value from large to small, denoted by vector **z**, where **z**_**j**_ = *k*, if the *j*th largest expression value in 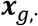 happens to be of cell type *k*. Denote ***O***_*k*_ as the cumulative frequency of cell type *k* over the expression order of 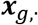, which is calculated as:

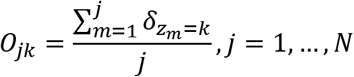

where 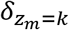 is the indicating function for *z*_*m*_ = *k*. A discrepancy score vector ***d*** between the observed and expected cell type frequency was further defined as

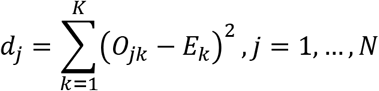

where ***d*** is a non-negative vector of length *N*, and it attains a minimum value of zero at *N*. The larger the maximum value *d* suggests the expression values are more enriched in certain cell types than the others. Denote *m* as the index of the maximum of *d*, i.e. *d*_*m*_ = max (*d*_*j*_), and the cell type frequency at the best discrepancy as 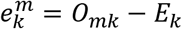, the cell types were further ordered by 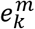 from large to small and denoted as ***o***, where ***o***_*j*_ = *k* if the *j*th largest value of 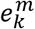 happens to be of cell type *k*. Then for cell type ***o***_1_ to ***o***_***K***_, the labeling matrix was built by 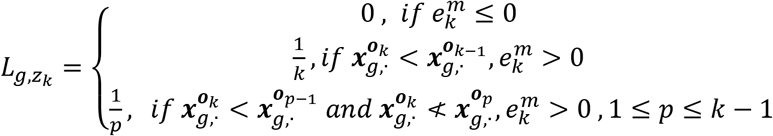, where 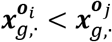 denotes *g* is significant over expressed in cell type ***o***_*j*_ compare to cell type ***o***_*i*_, which is tested by Mann Whitney test.

### Exclusion of the expression of undesired cells

ICTD can eliminate the expression signal from undesired cell types to excluder those cells from further analysis. To do this, ICTD first identifies gene co-expression modules from the decentralized expression matrix of their marker genes by using WGCNA and computes the first row base of each module by using SVD ^25^. Then for each gene that is positively co-expressed with one or several module(s) of the undesired cell type, its expression are further projected to the complementary space spanned by the first row base of each of such modules (s). Denote a decentralized tissue data as X, the data of pseudo-code of exclusion of the expression of undesired cells are given below:

**Algorithm 2:**
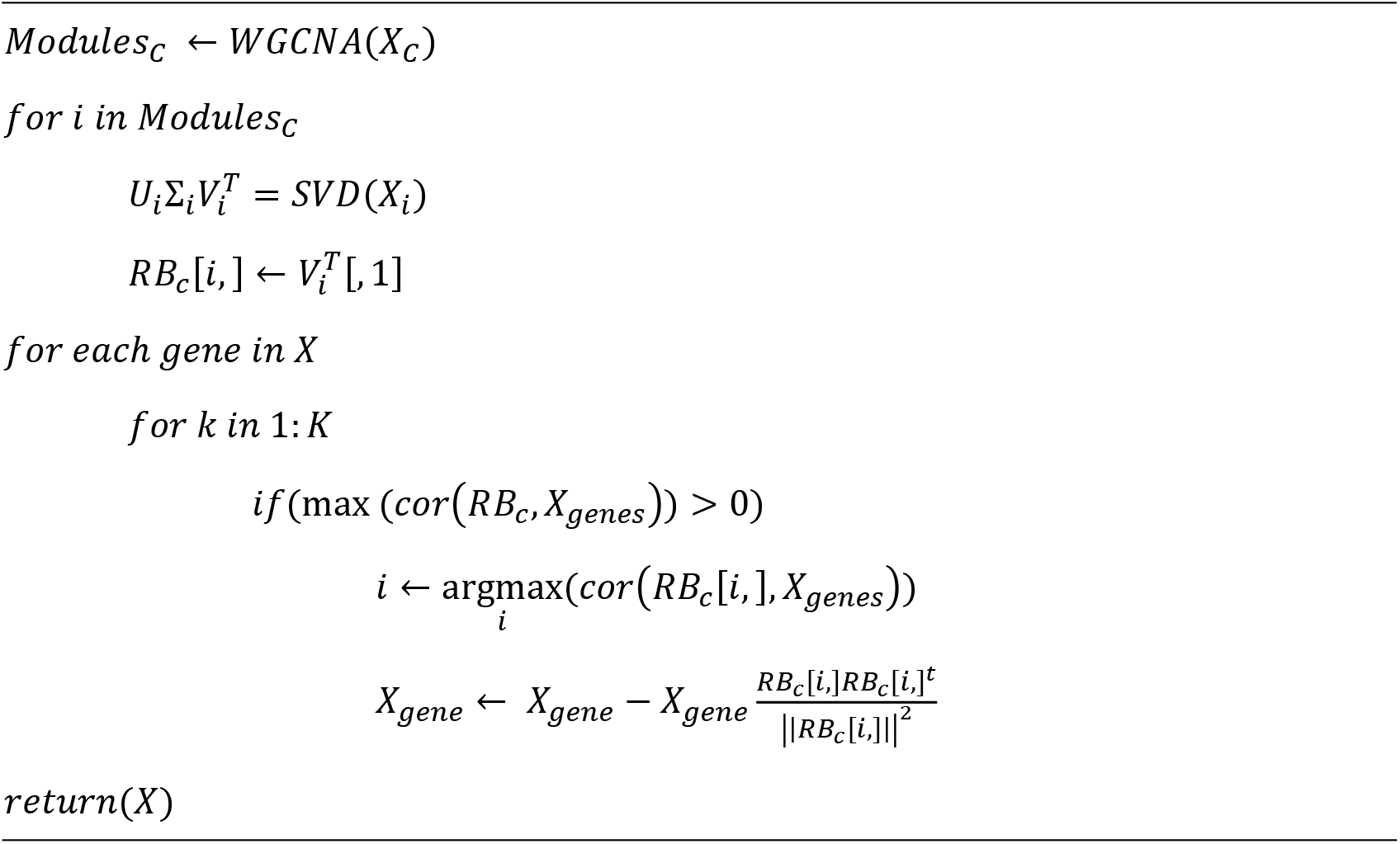
Remove the low rank space of undesired cell types

In this paper, we first identified 1089 cancer cell genes, as evidenced by their consistent up-regulation in 11 cancer types of TCGA data and significant expression in CCLE cell line data (Supplementary Table S10). Differential gene expression analysis was conduct by using Mann-Whitney test with FDR<0.05 as the significant cutoff and significant expression in cancer cell line data is determined by log(FPKM)>2. In the analysis of one specific cancer type, gene co-expression modules of the cancer genes were first identified. The linear space spanned by the modules were further excluded by the complementary space projection. Our analysis on single cell simulated and real bulk tissue data validated that such an elimination procedure can largely remove the expression of the genes stably expressed in cancer cells while retaining the low rank structure of the gene expressions from other cells (See **Supplementary Notes**).

### ICTD Step 2: Identification of rank-1 modules

Highly co-expressed modules were identified using our in-house method, namely MRHCA ^26 27^. More details about the MRHCA based module identification and its rationality in our case are given in **Supplementary Methods.**

The BCV test described in **Algorithm 1** is further applied to find the modules of rank-1, which possibly correspond to marker genes of identifiable cell types. The matrix rank of a module centered by a cell type uniquely expressed genes always increases with the module size, due to the genes less co-expressed with the hub may be expressed by other cell types. In this paper, we selected the modules of with hub significance p<1e-3, average co-expression correlation>0.8, rank=1 (p<1e-3) and with at least seven genes, as possible markers of identifiable cell types.

### ICTD Step 3: Determine the number and select Rank-1 modules of “identifiable” cell types

After identifying all sets of rank-1 marker genes, ICTD further determines the number of identifiable cell types, eliminates redundant and insignificant cell type marker genes, annotates each set of marker genes with a most likely cell type by using the labeling matrix, and build a marker gene – cell type representing matrix for the downstream deconvolution analysis.

Denote a rank-1 marker set 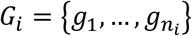 and labeling matrix *L*_*M*×*K*_, we first compute *S*_*i*_ = {*s*_*i*,1_, …, *s*_*i*,*K*_}, where 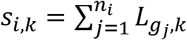 representing the enrichment level of *G*_*i*_ to the genes top expressed in cell type k. The significance level of *s*_*i*,*k*_, 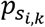, is assessed by a permutation test, and *G*_*i*_ is annotated as cell type with the minimal 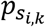 if 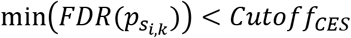. In this study, *Cutoff*_*CES*_ is selected as 0.01. The rank-1 markers annotated without a significant cell type annotation are excluded from further analysis. It is noteworthy that a larger *Cutoff*_*CES*_ can be selected for identification of possible unknown cell types.

Rather than predefining the cell types, ICTD determines the cell types that are “identifiable”. In some circumstance, the proportion of the cell type with a lower resolution is a non-negative linear sum of the proportion of several cell types with higher resolutions, such as the myeloid cell proportion equals to the sum of macrophage and neutrophils when these two cell types dominate the myeloid cell populations in the tissue ^28^. This linear dependency may correspond to a linear dependency between the row base of marker genes of cell types of different resolutions, which may result in number of identifiable cell types exceeding the rank of the linear space generated by the identified rank-1 markers.

To determine the number of identifiable cell types covered by the rank-1 marker genes, ICTD first construct a tree structure to represent the linear dependency among the identified rank-1 marker sets. A rank-1 marker set is considered as a root node if its row base can be non-negatively fitted by the row bases of other nodes with 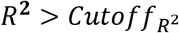. In this study, 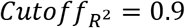 is selected. The rank-1 marker sets fitting each other with 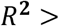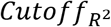 are merged together. All the root rank-1 marker sets are considered as markers of “identifiable” cell types and excluded from the further analysis. ICTD further computes the rank of the expression matrix of all the non-root rank-1 maker genes. Denoting the number of non-root rank-1 maker sets and their total rank as *P* and 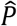. The total number of “identifiable” cell types among the non-root rank-1 marker sets is determined as 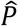.

A marker gene – cell type representation matrix is further computed for the downstream NMF analysis. Denote a selected rank-1 marker set as 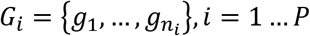, its gene expression profile as 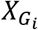, and ot SVD as 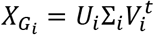, 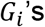 self-explanation score is defined as 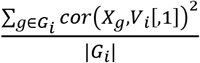, i.e. the averaged R square of the genes’ expression fitted by their first row base. The marker gene – cell type representation matrix C is constructed by **Algorithm 3**:

**Algorithm 3:**
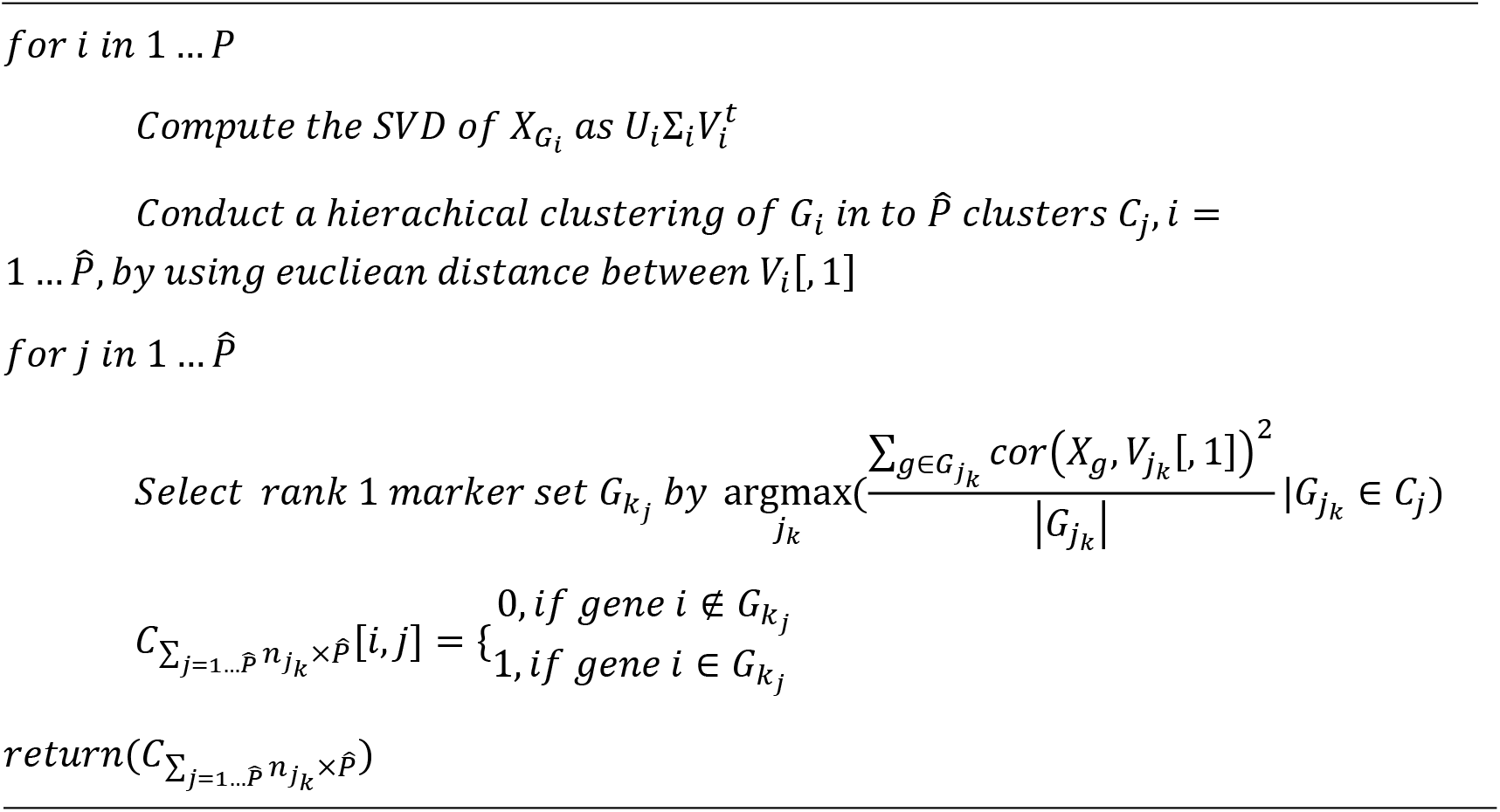
Construction of representation matrix

This step assigns marker genes of identifiable cell types that highly determines the prediction accuracy of the deconvolution analysis. ICTD also includes three other options in constructing marker genes and C matrix of identifiable cell types. The computational details and performance comparison of these methods were given in **Supplementary Methods.**

### ICTD Step 4: Constrained Non-negative Matrix Factorization

With the NMF constraint matrix 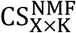, each of the K cell type is assigned with at least one cell type uniquely expressed gene (see derivations in method), hence the constraint NMF problem X_M×N_ = S_M×K_ · P_K×N_, S[I, k] ≥ 0, P[k, j] ≥ 0, S[I, k] = 0 *if* CS^NMF^[I, k] = 0 does have a unique solution ^29^. The rationale here is that the analysis only focuses on cell types with uniquely expressed markers that form rank-1 structure, and the analysis is robust to collinearity of cell proportions due to the uniqueness of solution. Specifically, for the *p*th disconnected subgraph with M_p_ genes, rank= K_p_, and constraint matrix C_*M*×K_, the NMF of X_*M*×N_ = S_*M*×K_ · P_K×N_ is solved by 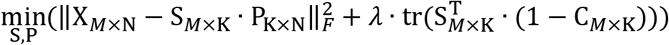, where S_*M*×K_ and P_*M*×K_ are the predicted signature and proportion of K cell types. Variables with fitted S that are highly varied from C are further removed. Detailed solution of the constrained NMF problem was given in **Supplementary Methods**. It is noteworthy when *λ* → ∞, *P*_*i*,*j*_ is the first row base of the SVD of 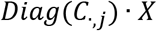, where 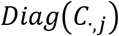 is the diagonal matrix generated by 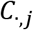. In this study, λ was selected based the best prediction accuracy trained on single cell simulated bulk data.

### ICTD Step 5: Co-deconvolution of matched multi-omics data

Multi-omics, including epigenetic and chromatin profiles, provide equally important characterization of tissue compositions as transcriptomic profiles. When multiple omics data are available for the same tissues, it is reasonable to assume that cell relative proportions deduced from each of the omics profile should be strongly associated. Based on this, co-deconvolution of matched multi-omics data could be used to cross-validate and robustify the proportion predictions, as detailed in **Algorithm 4**:

**Algorithm 4:**
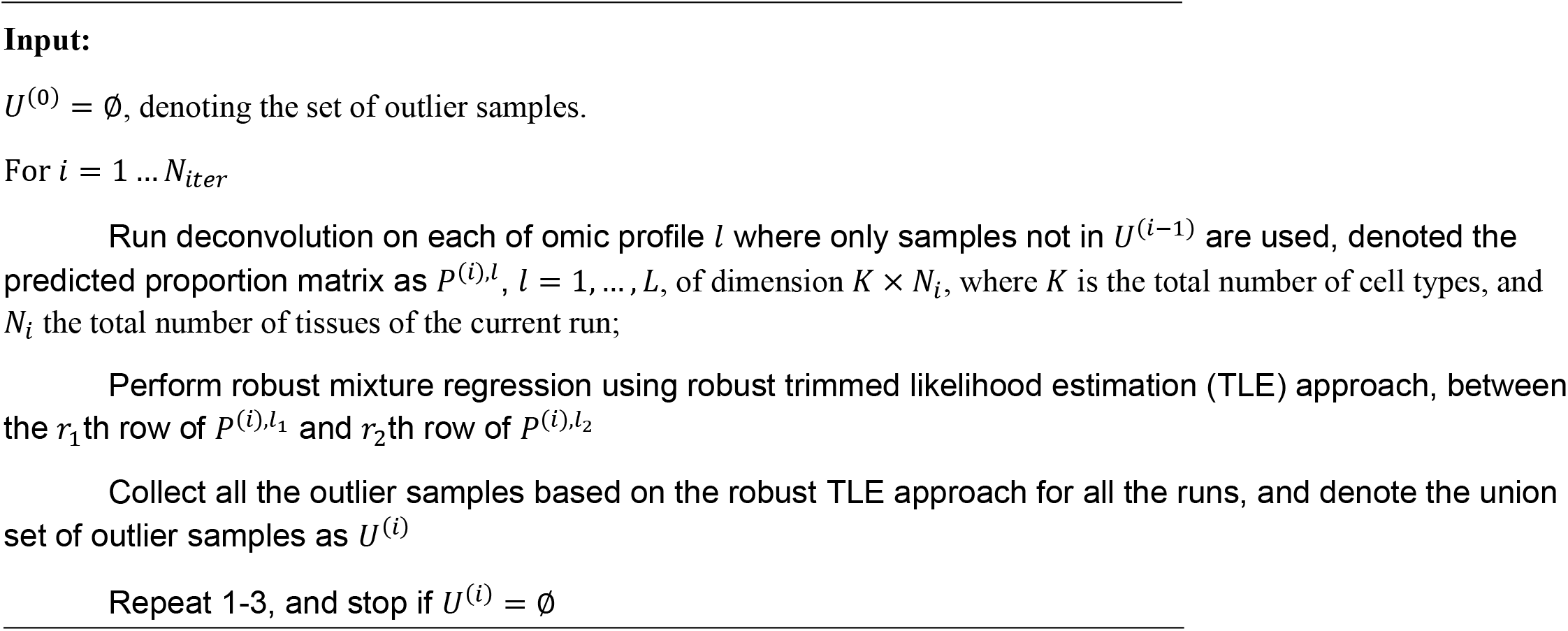
Co-deconvolution of matched multi-omics data

### ICTD Step 6: Conditional local low rank test of cell type varied function

Identifiable cell type specific function is defined by a group of genes that form a local rank-1 structure conditional on the estimated proportion of the cell type. A kernal function based local low rank structure screening method is developed for identification of such local rank-1 structures. Denote 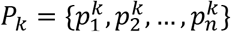 as predicted proportion of cell type k through the n samples and *P*_(*k*)_ = {*p*_*k*(1)_, …, *p*_*k*(*n*)_} as sorted *P*_*k*_ with an increasing order, 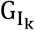 as the rank-1 marker genes of cell type k, and 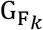 is a gene set containing possible marker genes of a varied function of k, the level of functional activity and its associated marker genes can be identified by **Algorithm 5**:

**Algorithm 5:**
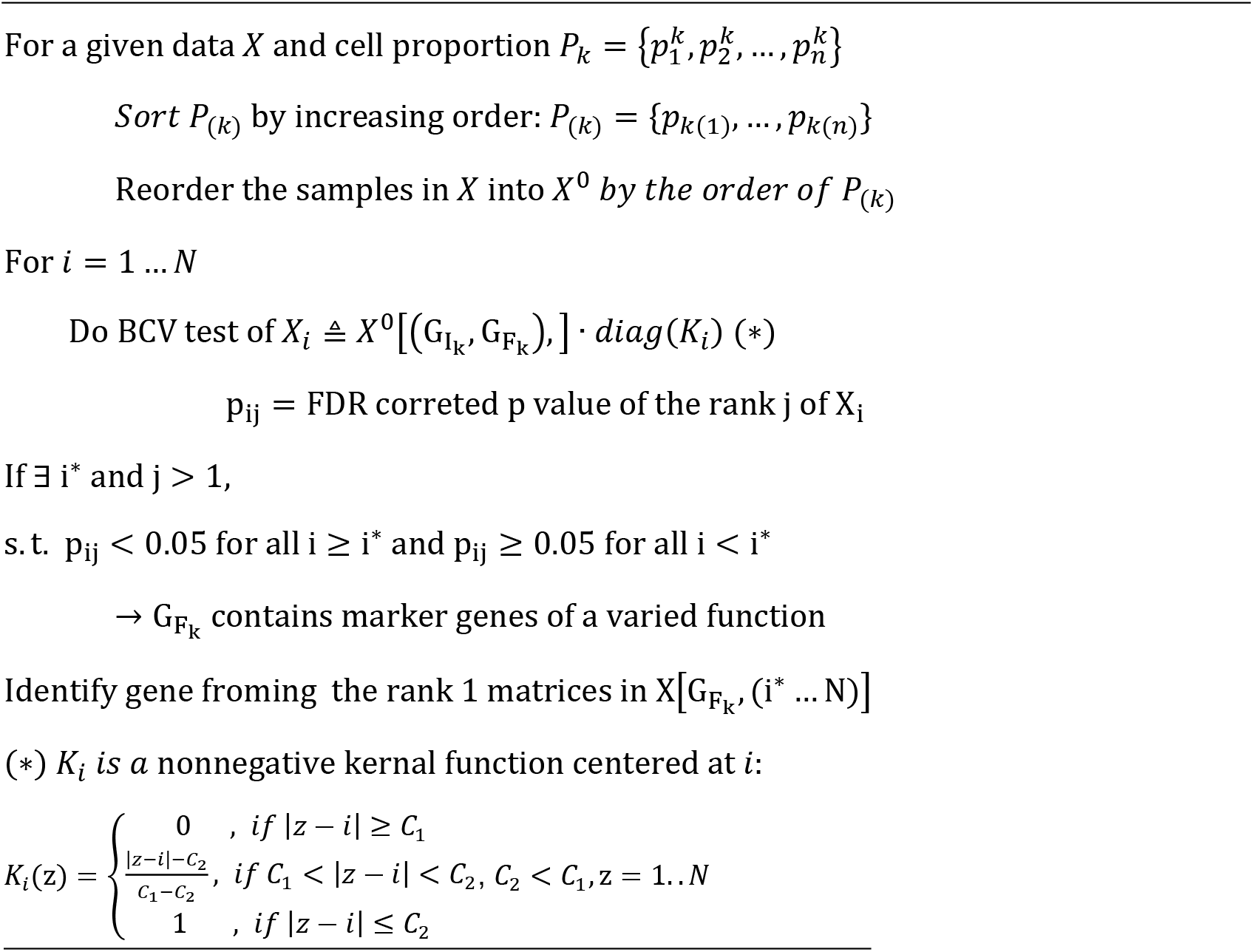
BCV screening of a local low rank structure

The idea of this algorithm is that the genes of a cell type specific function may form additional ranks in the samples with high proportion of the cells, which can be identified by the BCV test when only looking at those samples. The kernel function is to smooth the inter-sample variation in cell proportions (see more details in **Supplementary Methods**).

In this paper, 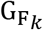 is selected for each cell type k by the genes annotated as top expressed by cell type *k* in the labeling matrix and with more than 0.8 co-expression correlation with the cell type *k*’s proportion. ICTD enables users to predefine 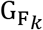 and select proportion of cell type *k* for a specified analysis, such as using known markers for prediction of T cell cytotoxicity ^30^. The functional activity level of each set of gene markers are then predicted by its first row base in the samples i ≥ i* by SVD. Averaged activity level per cell is further estimated by dividing the predicted functional activity level by the predicted cell type proportion.

### Single cell simulated Bulk Tissue data

Cell types in each scRNA-seq data were labeled by the cell clusters provided in the original works or by using Seurat pipeline with default parameters. Detailed information of the scRNA-seq data and cell type annotation is given in **Supplementary Table S3 and Notes**. For each data set, we simulate bulk tissue data with three steps: randomly generate the proportion of each cell type, called true proportion in this paper, that follows a Dirichlet distribution, (2) enforce a certain co-infiltration level of two selected cell types, and (3) draw cells randomly from the cell pool with replacement according to the cell type proportion, and sum up the expression values of all cells to produce a pseudo bulk tissue data. More details are provided in **Algorithm 6**:

**Algorithm 6:**
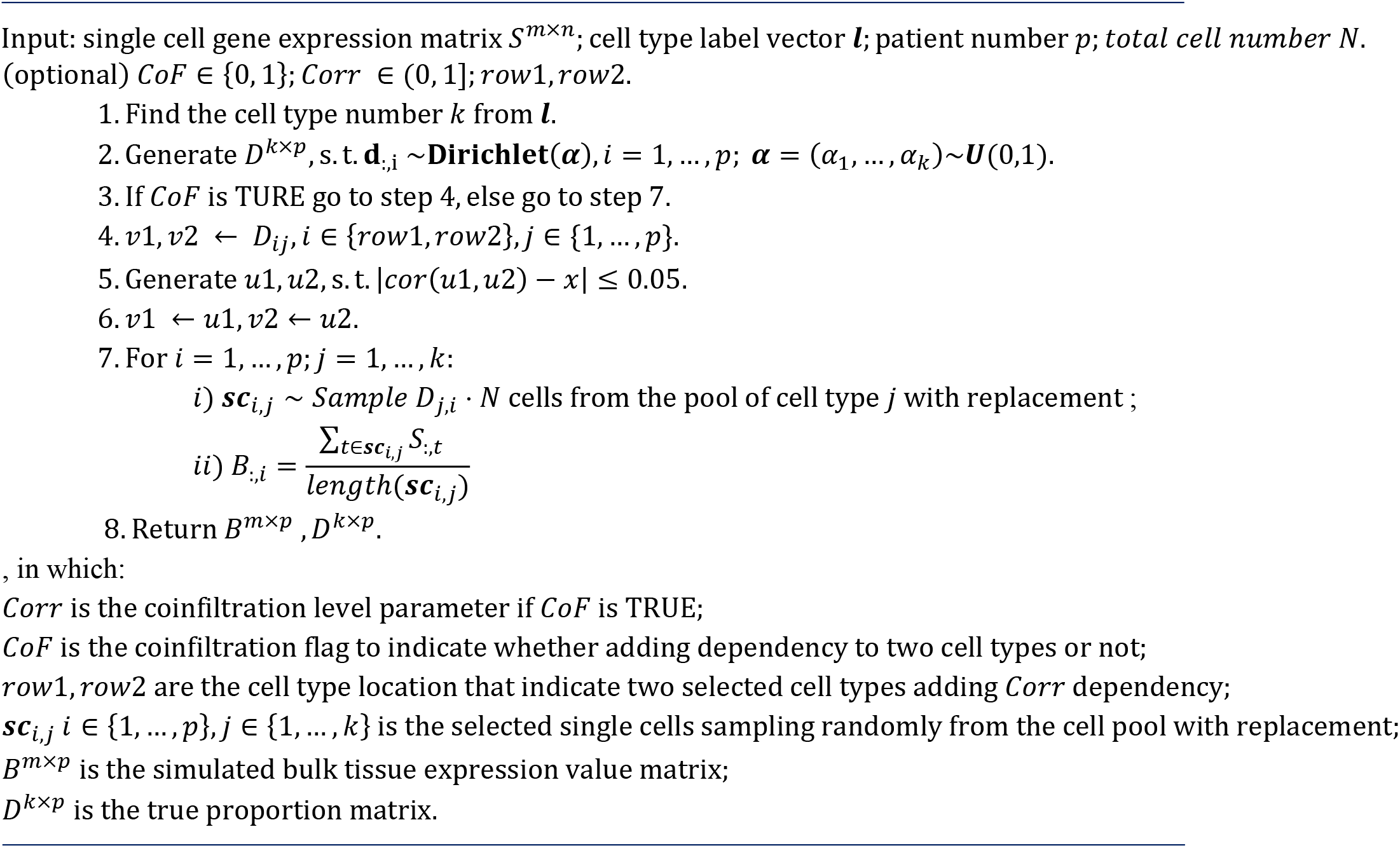
simulate bulk data using single cell

The Dirichlet distribution matrix was generated with R package “DirichletReg” (version 3.5.3). In order to evaluate the robustness of the deconvolution method while co-infiltration exists, we add different levels of co-infiltrations in our simulated bulk data to four pairs of cells that are commonly known to co-infiltrate in cancer tissue, namely, B/T cell, T/NK cell, Fibroblast/Endothelial cell, and B/Dendritic cell. (Supplementary Figure S13). For a robust method evaluation, five replicates were generated in the simulation of each data set and at each co-infiltration parameter.

### Explanation Score to evaluate the performance of our deconvolution method

We assessed the methods’ performance by the correlation between predicted and known proportion of each cell type in simulated data, which is inapplicable in the real tissue data. Thus, we developed

An explanation score (ES) was developed to evaluate the goodness that each marker gene’s expression is fitted by the predicted cell proportions:

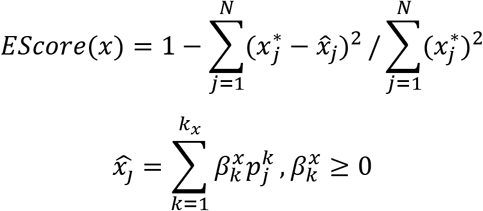

where 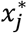 is the observed expression of marker gene *x* in sample *j*, 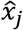 is the *x*’s expression level in *j* predicted by a non-negative regression model of the predicted proportion 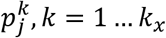 of *k*_*x*_ cell types that express *x*, and 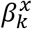 are parameters. Intuitively, with correctly selected marker genes, the marker gene’s expression can be well explained by the predicted proportions of the cell types that express the gene. Hence, a high ES score is a necessary but not sufficient condition for correctly selected marker genes and predicted cell proportion.

